# Is chaos making a difference? Synchronization transitions in chaotic and nonchaotic neuronal networks

**DOI:** 10.1101/224451

**Authors:** Kesheng Xu, Jean Paul Maidana, Samy Castro, Patricio Orio

**Affiliations:** Centro Interdisciplinario de Neurociencia de Valparaíso, Universidad de Valparaíso, Valparaíso, Chile; Instituto de Neurociencia, Facultad de Ciencias, Universidad de Valparaíso, Valparaíso, Chile

## Abstract

Chaotic dynamics of neural oscillations has been shown at the single neuron and network levels, both in experimental data and numerical simulations. Theoretical studies over the last twenty years have demonstrated an underlying role of chaos in neural systems. Nevertheless, whether chaotic neural oscillators make a significant contribution to relevant network behavior and whether the dynamical richness of neural networks are sensitive to the dynamics of isolated neurons, still remain open questions. We investigated transition dynamics of a medium-sized heterogeneous neural network of neurons connected by electrical coupling in a small world topology. We make use of an oscillatory neuron model (HB+I_h_) that exhibits either chaotic or non-chaotic behavior at different combinations of conductance parameters. Measuring order parameter as a measure of synchrony, we find that the heterogeneity of firing rate and types of firing patterns make a greater contribution than chaos to the steepness of synchronization transition curve. We also show that chaotic dynamics of the isolated neurons do not always make a visible difference in process of network synchronization transitions. Moreover, the macroscopic chaos is observed regardless of the dynamics nature of the neurons. However, performing a Functional Connectivity Dynamics analysis, we show that chaotic nodes can promote what is known as the multi-stable behavior, where the network dynamically switches between a number of different semi-synchronized, metastable states.

## Introduction

Over the past decades, a number of observations of chaos have been reported in the analysis of time series from a variety of neural systems, ranging from the simplest to the more complex [1, 2]. It is generally accepted that the inherent instability of chaos in nonlinear systems dynamics, facilitates the extraordinary ability of neural systems to respond quickly to changes in their external inputs [3], to make transitions from one pattern of behavior to another when the environment is altered [4], and to create a rich variety of patterns endowing neuronal circuits with remarkable computational capabilities [5]. These features are all suggestive of an underlying role of chaos in neural systems (For reviews, see [5–7]), however these ideas may have not been put to test thoroughly.

Chaotic dynamics in neural networks can emerge in a variety of ways, including intrinsic mechanisms within individual neurons [8–12] or by interactions between neurons [3,13–21]. The first type of chaotic dynamics in neural systems is typically accompanied by microscopic chaotic dynamics at the level of individual oscillators. The presence of this chaos has been observed in networks of Hindmarsh-Rose neurons [8] and biophysical conductance-based neurons [9–12]. The second type of chaotic firing pattern is the synchronous chaos. Synchronous chaos has been demonstrated in networks of both biophysical and non-biophysical neurons [3,13,15,17,22–24], where neurons display synchronous chaotic firing-rate fluctuations. The latter cases, the chaotic behavior is a result of network connectivity, since isolated neurons do not display chaotic dynamics or burst firing. More recently, a totally different mechanism showed that asynchronous chaos, where neurons exhibit asynchronous chaotic firing-rate fluctuations, emerge generically from balanced networks with multiple time scales synaptic dynamics [20].

Different modeling approaches have been used to uncover important conditions for observing these types of chaotic behavior (in particular, synchronous and asynchronous chaos) in neural networks, such as the synaptic strength [25–27] in a network, heterogeneity of the numbers of synapses and their synaptic strengths [28, 29], and lately the balance of excitation and inhibition [21]. The results obtained by Sompolinsky et al. [25] showed that, when the synaptic strength is increased, neural networks displays a highly heterogeneous chaotic state via a transition from an inactive state. Some other studies demonstrated that chaotic behavior emerges in the presence of weak and strong heterogeneities, for example a coupled heterogeneous population of neural oscillators and the difference of their synaptic strengths [28–30]. Recently, Kadmon et al. [21] highlighted the importance of balance of excitation and inhibition on a transition to chaos in random neural networks. All these approaches identify the essential mechanisms for generating chaos in neural networks. However, they give little insight into whether chaotic neural oscillators make a significant contribution to relevant network behavior, such as synchronization. In other words, whether the dynamical richness of neural networks are sensitive to the dynamics of isolated neurons has not been systematically studied yet.

To cope with this question, in the present paper we studied synchronization transition in heterogeneous networks of interacting neurons. Here we make use of an oscillatory neuron model (HB+I_h_) that exhibits either chaotic or non-chaotic behavior depending on biologically plausible parameter regions, as we showed in our previous study [12]. We simulated small-world [31] neural networks consisting on a heterogeneous population of HB+Ih neurons, connected by electrical synapses, and sampling their parameters from either chaotic or non-chaotic regions of the parameter space.

Our first finding is that isolated chaotic neurons in networks do not always make a visible difference in process of network synchronization transitions. The heterogeneity of firing rate and the types of firing patterns make a greater contribution to the steepness of the synchronization transition curve. Moreover, the macroscopic chaos is observed regardless of the dynamics nature of the neurons. However, the results of Functional Connectivity Dynamics (FCD) analysis show that chaotic nodes can promote what is known as the multi-stable behavior, where the network dynamically switches between a number of different semi-synchronized, metastable states. Finally, our results suggest that chaotic dynamics of the isolated neurons is not a predictor of macroscopic chaos, which can be a predictor of meta and multistability.

## Materials and methods

### Single neuron dynamics

We use a parabolic bursting model inspired by the static firing patterns of cold thermoreceptors, in which a slow sub-threshold oscillation is driven by a combination of a persistent Sodium current (*I_sd_*), a Calcium-activated Potassium current (*I_sr_*) and a hyperpolarization-activated current (*I_h_*). Depending on the parameters, it exhibits a variety of firing patterns including irregular, tonic regular, bursting and chaotic firing [12, 32]. Based on the Huber & Braun (HB) thermoreceptor model [11], it will be referred to as the HB+I_h_ model. Examples of the chaotic and non-chaotic oscillations are plotted in Fig 1 at different combinations of conductance parameters.

**Fig 1.**
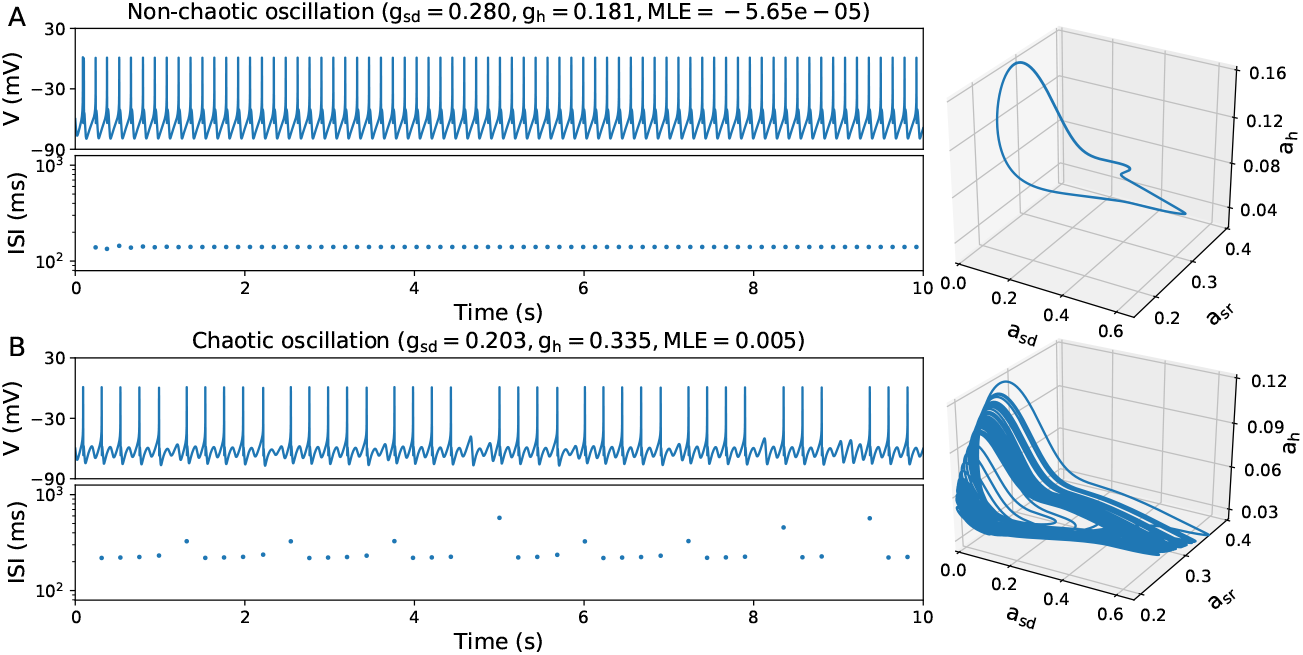
Examples of non-chaotic (A) and chaotic oscillation (B) of HB+I_h_ neurons at different combinations of conductance parameters. Below each voltage time course, inter-spike interval (ISI) plots are shown. Right panels show three-dimensional phase space projections of variables *a_sd_, a_sr_* and *a_h_*.

The membrane action potential of a HB+I_h_ neuron follows the dynamics:

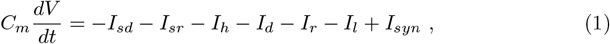

where *V* is the membrane capacitance; *I_d_, I_r_, I_sd_, I_sr_* are depolarizing (Na_V_), repolarizing (K_dr_), slow depolarizing (Na_P_ / Ca_T_) and slow repolarizing (K_Ca_) currents, respectively. *I_h_* stands for hyperpolarization-activated current, *I_l_* represents the leak current, and lastly the term *I_syn_* is the synaptic current. Currents (except *I_syn_*) are defined as:

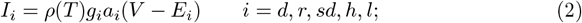

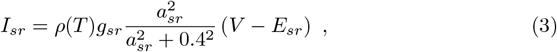

where *a_i_* is an activation term that represents the open probability of the channels (*a_l_* = 1), with the exception of *a_sr_* that represents intracellular Calcium concentration. Parameter *g_i_* is the maximal conductance density, *E_i_* is the reversal potential and the function *ρ*(*T*) is a temperature dependent scale factor for the current.

The activation terms *a_r_, a_sd_* and *a_h_* follow the differential equations:

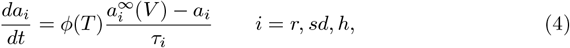

where

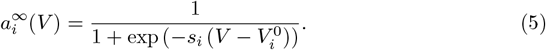

On the other hand, *a_sr_* follows

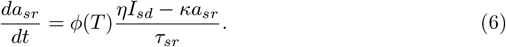

Finally,

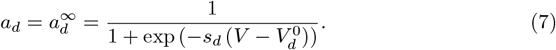

The function *φ(T)* is a temperature factor for channel kinetics. The temperature-dependent functions for conductance *ρ(T)* in Eq.(2)-(3), and for kinetics *φ(T)* in Eq.(4) and (6) are given, respectively, by:

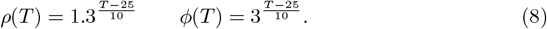

In the simulation, we vary the maximal conductance density *g_sd_, g_sr_* and *g_h_* values. Unless stated otherwise, the parameters used are given in Table 1.

**Table 1.**
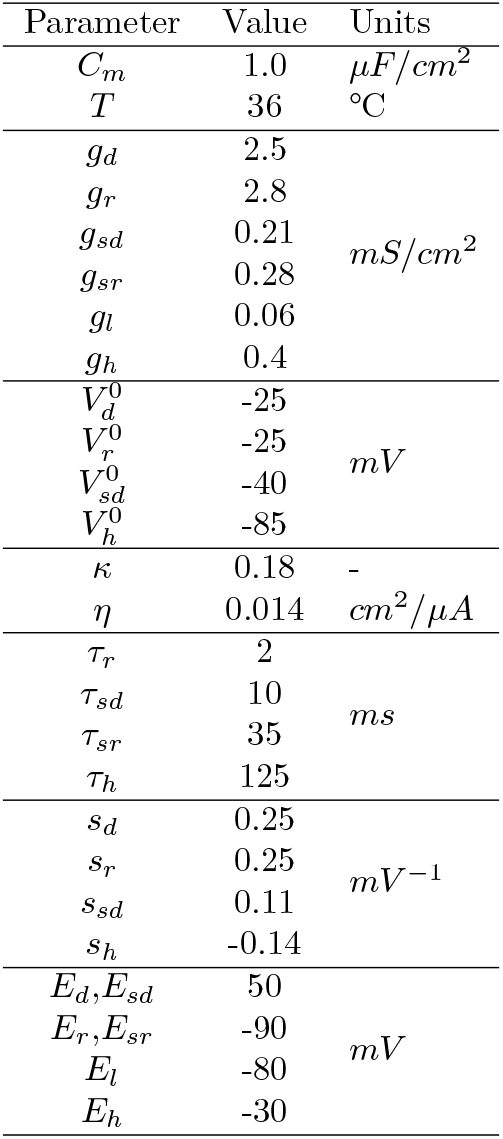
Parameters of the HB+I_h_ model.

### Synaptic interactions

The synaptic input current into neuron *k* is given by:

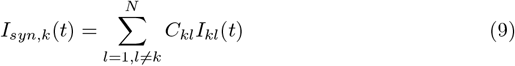

In this article, the current *I_kl_* between neuron *k* and *l* is modeled as a gap junction (electrical synapse):

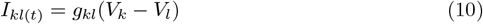

Where *g_kl_* is the conductance (coupling strength) of synapse from cell *k* to cell l. In this work we selected a uniform value *g_kl_ = g* for all connections within the neural networks, and simulations were performed with different values of *g* in order to observe the transition to total network synchrony.

### Structural connectivity matrix

We define the connectivity matrix by *C_kl_* = 1 if the neuron *k* is connected to neuron *l* and *C_kl_* = 0 otherwise. We employed a Newman-Watts small world topology [31], implemented as two basic steps of the standard algorithm [33–35]: (1). create a ring lattice with *N* nodes of mean degree 2*K*. Each node is connected to its *K* nearest neighbors on either side. (2). For each node in the graph, add an extra edge with probability *p*, to a randomly selected node. The added edge cannot be a duplicate or self-loop. Finally, as we are simulating electrical synapses, the matrices were made symmetric. In our simulations, *N* = 50, *K* =1, *p* = 0.4. The random seed for adding extra edges was controlled in order to use the same set of connectivity matrices under each condition.

### Quantifying chaos

The method for establishing whether a system is chaotic or not is to use the Lyapunov exponents. In particular, Maximal Lyapunov exponent (MLE) greater than zero, hence, is widely used as an indicator of chaos [36–39]. We calculated MLEs from trajectories in the full variable space, following a standard numerical method based on that of Sprott (2003) [36] (see also Jones et al., 2009) [37].

### Measurement of network dynamics

The voltage trajectory of each neuron was low-pass filtered (50Hz) and a continuous Wavelet transform [40–42] was applied to determine in one step the predominant frequency and the instantaneous phase at that frequency. We use the complex Morlet wavelet as mother wavelet function to calculate instantaneous phase.

We describe global dynamical behavior of the neural networks using the mean and the standard deviation of the order parameter amplitude over a time-course, which indicate respectively the global synchrony and the global metastability of the system [43, 44]. The order parameter [45, 46], *R*, describes the global level of phase synchrony in a system of *N* oscillators, given by:

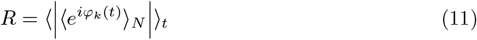

Where *φ_k_(t)* is the phase of oscillator *k* at time *t*, 〈*f*〉_*N*_ = *ϕ_c_(t)* denotes the average of *f* over all *k* in networks, | | is absolute value and 〈*f*〉_*t*_ is the average in time. *R* = 0 corresponds to the maximally asynchronous (disordered) state, whereas *R* = 1 represents the state where all oscillators are completely synchronized (phase synchrony state). The global metastability *χ* of neural networks is given by:

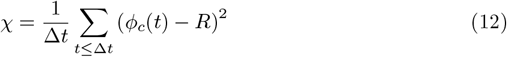

Metastability is zero if the system is either completely synchronized or completely desynchronized during the full simulation–a high value is present only when periods of coherence alternate with periods of incoherence [43], where Δ*t* of Eq.(12) is time windows to quantify the global metastability.

### Functional Connectivity Dynamics

A functional connectivity (FC) matrix was calculated by the pair-wise phase synchrony (order parameter). This was done in a series of *M* overlapping time windows *T*_1_, *T*_2_, *T*_3_, …, *T_M_*. The FC matrix at the window *m* is defined by:

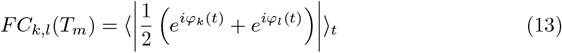

Where *t* corresponds to all times inside window *m*. The values in the lower triangle of FC, discarding the diagonal and the values adjacent to it, were vectorized and a correlation matrix is calculated between the vectors. This constitutes the Functional Connectivity Dynamics (FCD) matrix [47, 48]:

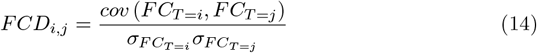

Where *cov* (*X,Y*) is the covariance between vectors *X* and *Y*, and *σ(X)* is the standard deviation of *X*. Note that the *(i,j)* indices in Eq.(14) refer to FC matrices obtained at different times, while the *(k,l)* indices in Eq.(13)refer to network nodes. Finally, an histogram of FCD values offers a rough measure of multistability (see Fig 6).

### Numerical integration

Equations (1)–(8) were solved by the Euler method with a fixed step size *dt* = 0.025. Most simulations were also repeated with an adaptive integration algorithm (odeint routine of Scipy package) without noticeable difference in the results. In our simulations we took *t* = 20000 time units after removing transient state for being in a stationary state. Data analysis and plotting were performed with Python and the libraries Numpy, Scipy, and Matplotlib.

## Results

### Synchronization transitions with parameters drawn from fixed-size regions

Our main goal is to study how the dynamics of isolated neural oscillators can propagate to the network level, in terms of relevant behaviors. To do this, we use a model of neural oscillator that can display either chaotic or non-chaotic behavior depending on the parameters (Fig 1, Fig 2A, Xu et al 2017 [12]). We simulated networks of 50 neurons connected by electrical coupling (gap junctions) in a small world topology. When drawing the *g_sd_* and *g_h_* parameters, we selected different regions of the parameter space that made them behave as either chaotic or non-chaotic oscillators, while maintaining a similar average firing rate (Fig 2A). Then, the inter-cellular conductance *g* was varied from 0 to 1 in order to evidence the transition from asynchrony to complete synchronization.

**Fig 2.**
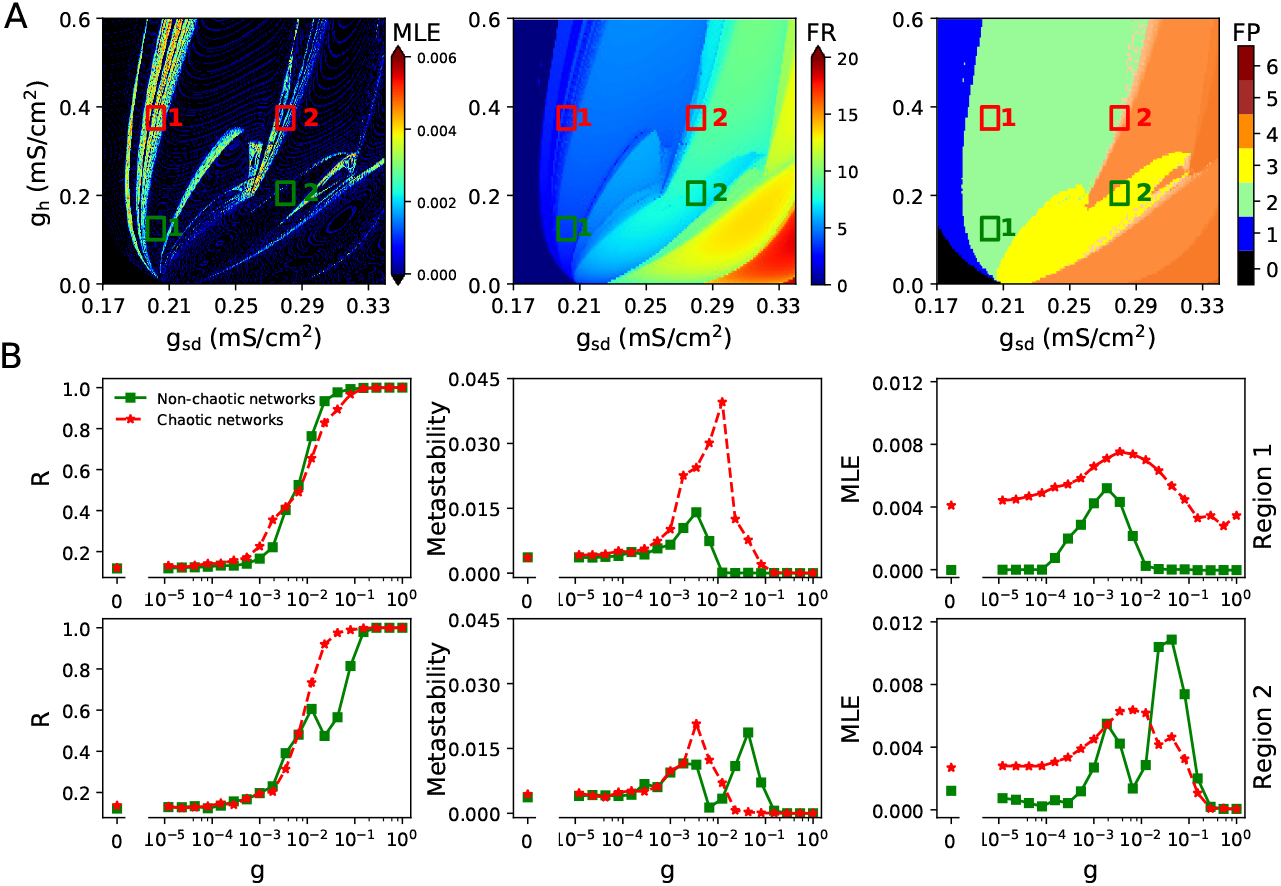
Synchronization transitions on the neural networks with parameters drawn from fixed-size regions of the parameter space. **A**. Maximal Lyapunov exponent (MLE), Firing rate (FR) and Firing pattern (FP) obtained from each parameter values. Regions marked in red correspond to chaotic oscillations, characterized by MLE>0, while regions in green are non-chaotic oscillators (MLE⩽0). The color bar of FP indicates: 0, no oscillations; 1, sub-threshold oscillations (no spikes); 2, oscillations and spikes with skipping; 3, regular tonic spiking; 4, burst firing (the shade represents the number of spikes per bursts); 5, tonic with firing rate between 20 and 50 spikes/second; 6, firing rate higher than 50 spikes/second. **B**. Transition dynamics in heterogeneous networks of 50 neurons, as the *g* coupling value is increased. Order parameter (R), metastability and MLE are shown. Throughout this article, Non-chaotic networks and Chaotic networks in legend respectively denote the networks built with non-chaotic and chaotic oscillators.

Fig 2B shows the transition curves for networks built with parameters drawn from regions 1 (top) and 2 (bottom), showing the order parameter (global synchrony), metastability (time variability of the global synchrony) and the network MLE as a measure of global chaos. In the first pair of parameter regions, we observe that networks of chaotic oscillators show a shallower transition to synchrony, with a higher metastability and a higher network MLE. This suggest that the chaotic nature of the oscillators indeed impacts the network behavior. In networks of non-chaotic oscillators, when *g* is between 10^-4^ and 10^-2^, we observe that the networks becomes chaotic while transition from asynchrony (low R) to synchrony (high R). However, when phase synchronization is reached, chaos is lost. Thus, only asynchronous chaos is observed. In the case of chaotic neurons, we find that the networks always exhibits chaotic behavior at a wide range (with *g* from 0 to 1) of synaptic coupling. In that case, asynchronous and synchronous chaos both are observed. When the same analysis was applied to the second pair of parameter regions, we observe a strange behavior of the non-chaotic oscillators, with a non-smooth transition associated to a higher metastability and network MLE. An inspection of the firing patterns (Fig 2A, right) reveals that the regions labeled as ‘2’ contains transitions between different firing patterns: tonic regular to bursting for non-chaotic, and skipping to bursting for the chaotic region. This made us think that the ‘kink’ observed in the curves of non-chaotic oscillators, was due to a transition between firing patterns occurring in the network. It can be seen in Fig 3A that as synaptic coupling is increased in networks of non-chaotic neurons, bursting firing pattern dissapear, while in chaotic neurons they are increased. Moreover, in non-chaotic neurons there is a rebound of bursting firing patterns at *g* = 0.0433. This finding suggests that the dramatical changes in firing patterns induce the non-smooth transition shown in Fig 2B, (bottom).

**Fig 3.**
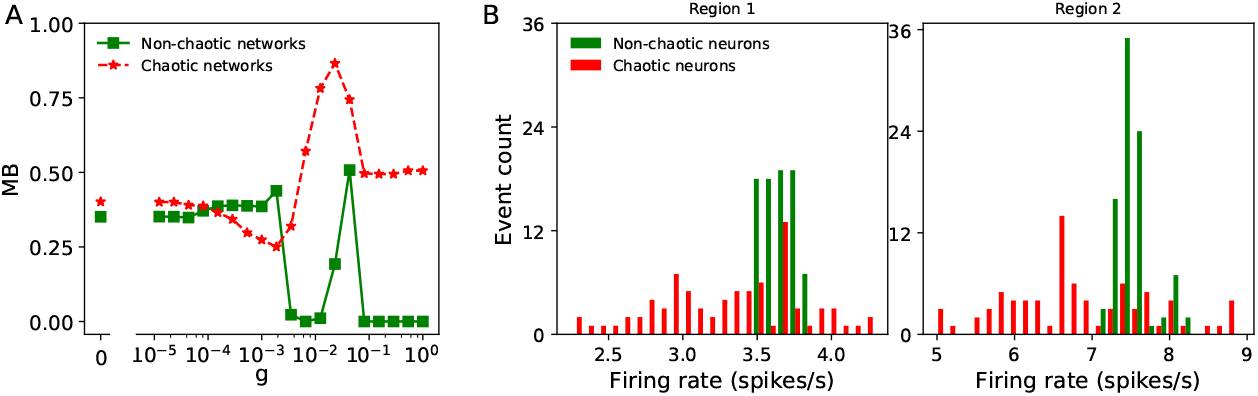
**A**. Mean fraction of bursting events (MB) in networks of chaotic and non-chaotic oscillators from region 2. The fraction of bursting events for a given neuron *k* is defined as: *b_k_* = *Nb/Te*, where *Nb* and *Te* present the number of bursters (two or more spikes) and total events for isolated neurons, respectively. Then, the mean fraction of bursting events of the whole neural networks is expressed as 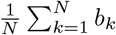. **B**. Histogram of the (isolated) firing rates in each pair of parameter regions that are shown in Fig 2A.

A closer examination of the isolated firing rates in each parameter range revealed another effect of chaos. A histogram of the (isolated) firing rates in each parameter region (Fig 3B) reveals that, when drawing from fix-sized regions of the parameter space, chaotic oscillators show a more heterogeneous distribution of firing rates than non-chaotic. Thus, the steeper transition to synchrony and lower metastability observed with non-chaotic oscillators could be due to a more homogeneous nature of the network, rather than the chaotic oscillation by itself. Although this is already an effect of chaos, it is of dubious biological relevance because neurons will never control their levels of channel expression within a fixed range, as we did here. If any, neurons control for function, and a simple approximation to this is to consider that they try to maintain a certain average firing rate with whatever ion channel density relationship that can attain it. Thus, we developed a parameter sampling procedure that replicated, for both chaotic and non-chaotic populations, a similar firing rate *distribution* rather than the mean. We also shifted to another parameter sub-space (*g_sd_/g_sr_*) to take advantage of the complete absence of chaos when *g_h_* = 0 (Xu et al [12]). Nevertheless, the simulations that follow were also performed in the *g_sd_/g_h_* parameter subspace with very similar results (see Supplemental Material S1)

### Synchronization transitions using same distribution of firing rates

Fig 4A shows a region of the *g_sr_/g_sd_* parameter sub-space, plotting Maximal Lyapunov exponent and firing rate obtained with each parameter pair. The example of desired regions with lower firing rate (from 3.0 to 4.5 spikes/s) is plotted in the firing rate (Fig 4A, middle and right). The regions shown in a darker tone of blue correspond, respectively, to chaotic (Fig 4A, middle) and non-chaotic oscillations (Fig 4A, right) behavior. The 3.0 - 4.5 Hz interval was divided in bins of 0.1 Hz, and in each bin the same number of *g_sr_/g_sd_* combinations was randomly picked from each region (chaotic or non chaotic). In addition, we picked parameter pairs from the model without the *I_h_* current (NoIh oscillators) that do not display chaotic behavior under any parameter combination (See Xu et al. 2017 [12] and Fig 4B). The desired region of this case in firing rate is shown in the Fig 4B (bottom). The histogram of firing rates in each selected set of parameters (Fig 4C), shows that they have the same distribution of firing rate. The same operation was used to select parameter sets with higher firing rate (from 7.0 to 9.5 Hz, not shown).

**Fig 4.**
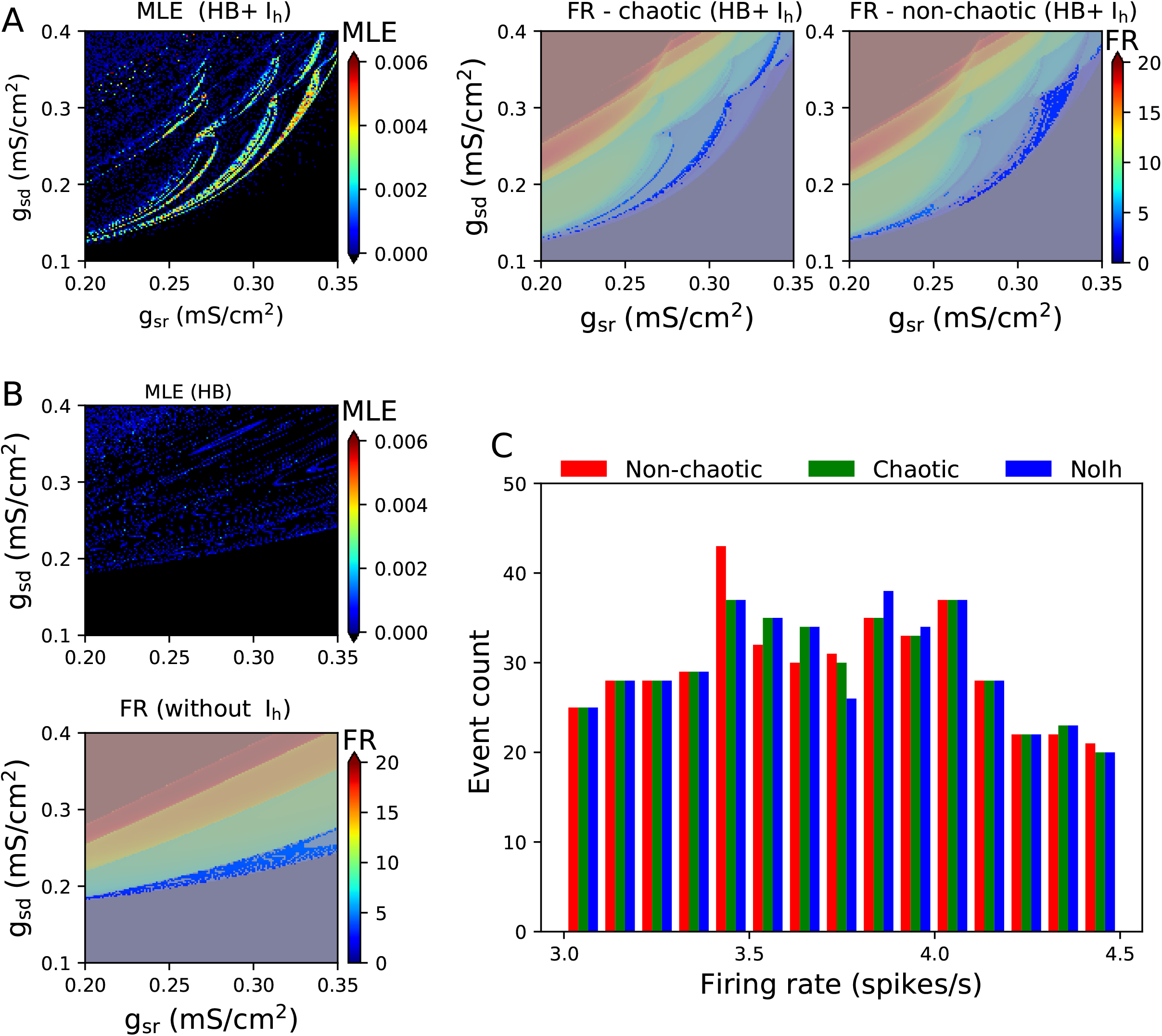
Selection of parameter sets with same distribution of firing rate. **A**. Maximum Lyapunov Exponent (MLE) and Firing Rate (FR) obtained in the selected *g_sr_/g_sd_* parameter region. The FR plot is shown twice, highlighted in light blue either chaotic (MLE > 0, left) or non-chaotic (MLE ⩽ 0, right) oscillations with a Firing Rate between 3.0 and 4.5 spikes/s. **B**. MLE and FR in the same *g_sr_/g_sd_* region as in **A**, for the model without *I_h_*. Highlighted in light blue, FR between 3.0 and 4.5 spikes/s. **C**. Histogram of firing rates in each selected set of parameters, showing the same distribution of firing rate

Networks of 50 neurons were built by randomly picking *g_sr_/g_sd_* pairs from the populations described above. Fig 5 plots the synchrony transition curves for networks using the parameters drawn from the lower (Fig 5A) and higher firing rate (Fig 5B) ranges, showing the order parameter, metastability and the network MLE. In the lower firing rate regions, all types of networks show a similar slope in their transition to synchrony. Networks of chaotic oscillators, however, show a higher metastability and a higher network MLE. In the higher firing rate regions, we observe that networks of both chaotic and non-chaotic oscillators show not only a similar transition to synchrony, but also the same degree of metastability and network MLE. These results suggest that MLE at the network level (macroscopic chaos) can be a predictor of metastability, however the chaotic nature of the isolated oscillators will not always translate to network chaos in a direct or predictable fashion. The blue curves of Fig 5 display that networks of NoIh oscillators show a transition to synchrony at lower values of *g*, with the similar degree of metastability and lower values of network MLE compared to the other networks. However, it is worth to mention that NoIh systems have 50 dimensions less (1 per node) and thus the magnitude of the Lyapunov exponents may not be comparable.

**Fig 5.**
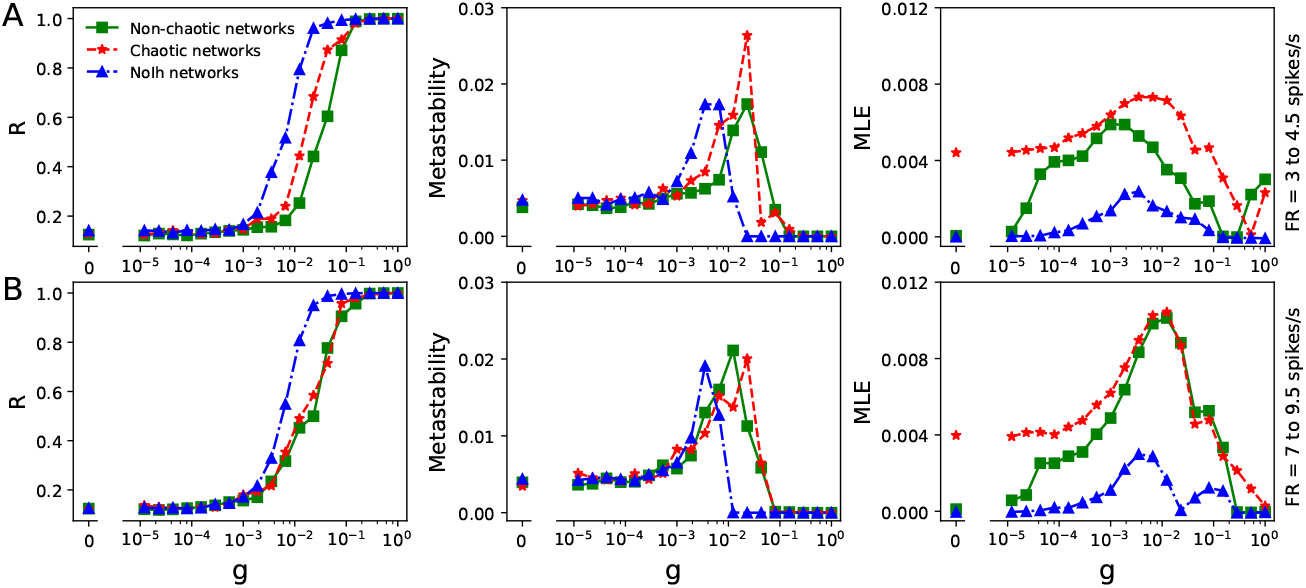
Transition curves for networks with parameters *g_sr_/g_sd_,* as described in Fig 4. Synchronization transition characterized by Order parameter, metastability and the network MLE. **A** and **B** denote parameters drawn from the lower and higher firing rate, respectively. Here NoIh networks in legend presents the network built with NoIh oscillators.

### Multi-stable behavior in neural networks

Finally, we measured the ability of our network models to display multi-stable behavior by characterizing their *functional connectivity dynamics* (FCD). This analysis is being extensively applied to fMRI and M/EEG recordings [47–49] and is explained in Fig 6 and Methods. Briefly, the series is divided in overlapping time windows and for each window a matrix of pair-wise synchrony between the nodes is calculated. Then, the synchrony matrices are compared against each other in the FCD matrix, where the axes now represent time.

**Fig 6.**
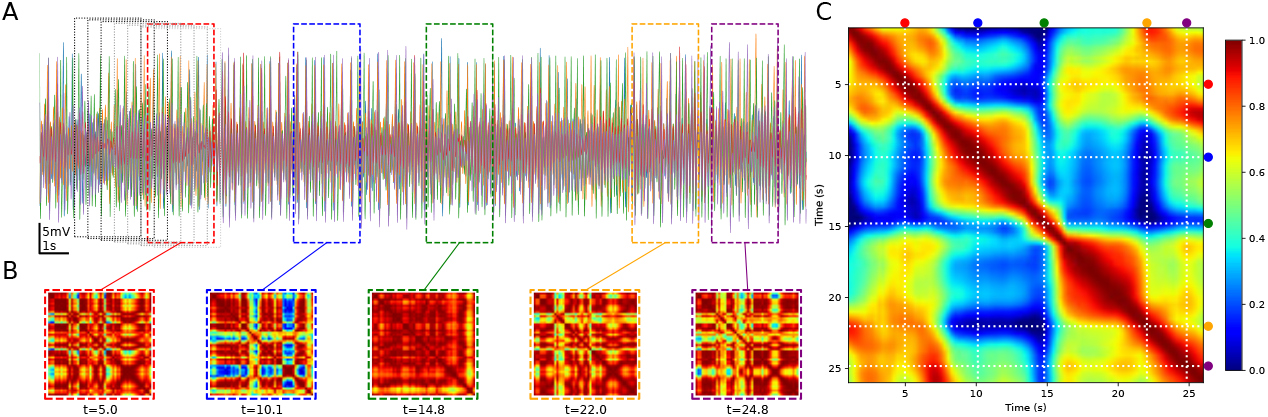
Functional Connectivity Dynamics Analysis. **A**. Time course of 50 chaotic nodes (only 5 traces are shown), showing the time windows for synchrony analysis. **B**. Functional Connectivity (FC) matrices obtained in 5 sample time windows. **C**. All the FCs are compared against each other by Pearson correlation and this constitutes the FCD matrix. The dotted lines and the color dots at right and top represent the FCs shown in B.

The FCD matrices that we obtained showed distinctive patterns for the unsynchronized and synchronized situations (Fig 7A). In the first case (synaptic conductance *g* = 0), all values outside the diagonal were 0 or close to 0. This means that the pair-wise synchronization patterns or FC matrices continuously evolve in time and never repeated during the simulation. On the other hand, when synaptic conductance is maximal, all the values in the matrix are equal to 1, meaning that the synchronization is the same and maintained through all the simulation. However, at intermediate values of *g,* some FCD matrices show a mixture of values between 0 and 1, with noticeable ‘patches’ that evidence the transient maintenance of some synchronization patterns. We call this a multi-stable regime. The histograms of FCD values (shown in Fig 7A below each FCD) are also useful in detecting the three situations described. As a rough measure of multi-stability, we took the variance of the FCD values (outside the diagonal) and plotted them against the synaptic conductance, averaging several simulations with different seed for the random connectivity matrix (Fig 7B). In the 3.0 to 4.5 firing rate range (for the isolated oscillators), it is clear that chaotic nodes produce networks with higher multi-stability than both non-chaotic and NoIh nodes. Moreover, the *g* range in which the multi-stable behavior is observed is wider. In the 7.0 to 9.5 firing rate range, the variance of the FCD is not higher for chaotic nodes, however the *g* range for multi-stability is still wider. This shows that FCDs with signatures of multi-stable behavior are more easily obtained when the networks are composed of chaotic nodes. FCD Analysis were also performed in the *g_sd_/g_h_* parameter subspace with very similar results shown in S2-Fig.

**Fig 7.**
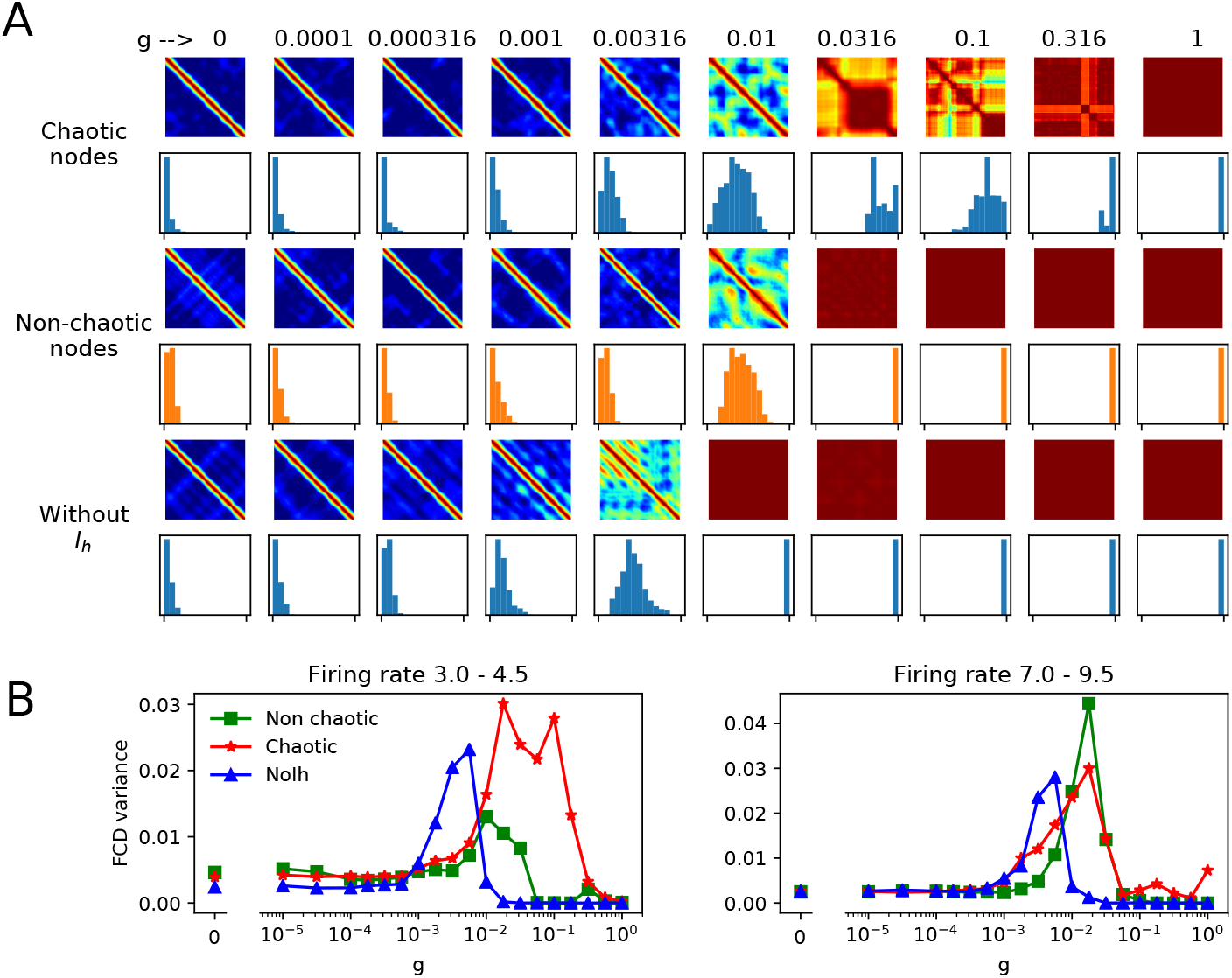
FCD in networks of chaotic and non-chaotic oscillators. **A**. FCD matrices obtained at different values of synaptic conductance *g* in networks of either chaotic, non-chaotic or NoIh nodes. Below each matrix, an histogram of the values is shown. The diagonal and the neighboring values were not included. **B**. Variance of the FCD values (the same values plotted in the histograms) plotted against *g*. Average of 15 simulations with different random seed for the small-world connectivity and parameters.

## Discussion

In this work, we investigated how a complex node dynamics can affect the synchronization behavior of a medium-sized heterogeneous neural network. We systematically controlled the chaotic nature of the oscillators, trying to keep other variables, such as network heterogeneity, constant. While some works have focused on network connectivity, only a few works have explored the impact of node dynamics to the network. Reyes et al. (2015) [50] showed that very small networks (2-3 neurons), when composed of irregular nodes can provide a wider frequency range. In medium-size networks but using a much simpler node dynamics, Hansen et al. (2015) [47] showed that bi-stable nodes enhance the dynamical repertoire of the network when looking at the FCD.

At first glance, our results are not as straightforward to interpret as in the previously mentioned works. Chaotic node oscillations do not always make a visible difference in terms of mean network synchronization and, most notably, chaos at the network level (macroscopic chaos) was always obtained regardless of the nature of the nodes. Other factors, such as network heterogeneity and the transitions between different firing patterns, seem to be more determinant to the steepness of the synchronization transition curves. However, we found that chaotic nodes can promote what is known as the multi-stable behavior, where the network dynamically switches between a number of different semi-synchronized, metastable states. Our results suggest that macroscopic chaos can be a predictor of metastability, as the greatest values of this measure (as well as multi-stability) coincide with intermediate *g* values where the maximum MLEs were found. However this must be taken with caution as the MLE is not necessarily a quantitative measure of chaos [51].

The chaotic nature of the isolated oscillators did not always convey to network chaos in a direct or predictable fashion. More specifically, our networks always showed chaotic behavior at some *g* values regardless of the dynamics of the isolated nodes. This is not surprising, as chaotic behavior arises in networks of very simple units and seems to depend more strongly on other factors such as synaptic weights and network topology [25-27, 52]. Moreover, just the high-dimensionality of the systems seems to be enough to assure that chaos will emerge under some conditions, for example, the quasiperiodic route to chaos in high-dimensional systems [53].

Assessing chaos in a large network or in a high-dimension system can be a difficult task. It is now generally accepted that a unique intrinsic and observable signature of systems exhibiting deterministic chaos is a fluctuating power spectrum with an exponential frequency dependency [54]. Thus, some studies introduced the broad power spectrum to characterize the chaos of networks [13, 55]. Here we use the most popular and directly method of maximal Lyapunov exponent (MLE) to quantify chaos on the level of networks in the way as we did for single cells [12, 36, 37]. As usual, we define the state of the network as chaotic if the MLE is greater than zero.

Macroscopic chaos mentioned above arises from the network’s global properties, the propensity of isolated neurons to oscillate, the nature of synaptic dynamics, or a mixture of the them, as shown in earlier works [25–29]. In this paper, the focus is different. First, finding both asynchronous and synchronous chaos in the same network, only by changing the synaptic strength, is new. Secondly, the route from asynchronous to synchronous chaos in networks of chaotic and non-chaotic oscillations has a slight difference and has not been found in previous studies. Specifically, network switches directly from asynchronous to synchronous chaos in networks of chaotic neurons, while networks of non-chaotic neurons usually can go through four phases of network state, that are asynchronous activity, asynchronous chaos, then again asynchronous activity and lastly synchronous chaos.

While discussing chaos in neural systems we have used completely deterministic dynamics. Random variables were used to define network connectivity and node parameters, but the time evolution of the networks and nodes was calculated in the absence of noise. However, neural systems are subject to a number of noise sources, being the most important the stochastic opening an closing of ion channels and synaptic variability [56]. How the synchronization transitions and meta/multi-stable behavior will emerge in a noisy system remains to be studied, and it will be interesting to assess how much the dynamics introduced by chaos can prevail in the presence of noise.

In summary, we have shown that chaotic neural oscillators can make a significant contribution to relevant network behavior, such as states transition and multi-stable behavior in neural networks.Our results open a new way in the study of the dynamical mechanisms and computational significance of the contribution of chaos in neuronal networks.

## Supporting information

S1-Fig. **Synchronization transitions on the case of** *g_h_/g_sd_* **with same distribution of Firing Rate.**

S1-Fig (A) shows FR (top) and MLE (bottom) plot obtained from each parameter values of *g_sd_/g_h_* parameter space. The regions shown in a darker tone of blue correspond, respectively, to chaotic (left) and non-chaotic oscillations (right) behavior. **S1-Fig (B)** plots the histogram of a population of (isolated) firing rates from each desired parameter regions, indicating the similar distribution of firing rate for chaotic and non-chaotic case. **S1-Fig (C)** plots the synchrony transition curves for networks using the parameters drawn from ranges 2 to 3.5 spikes/s, showing the order parameter, metastability and the network MLE. The results obtained here are compatible with what has been plotted of main text (shown in Fig 5).

**S1-Fig.**
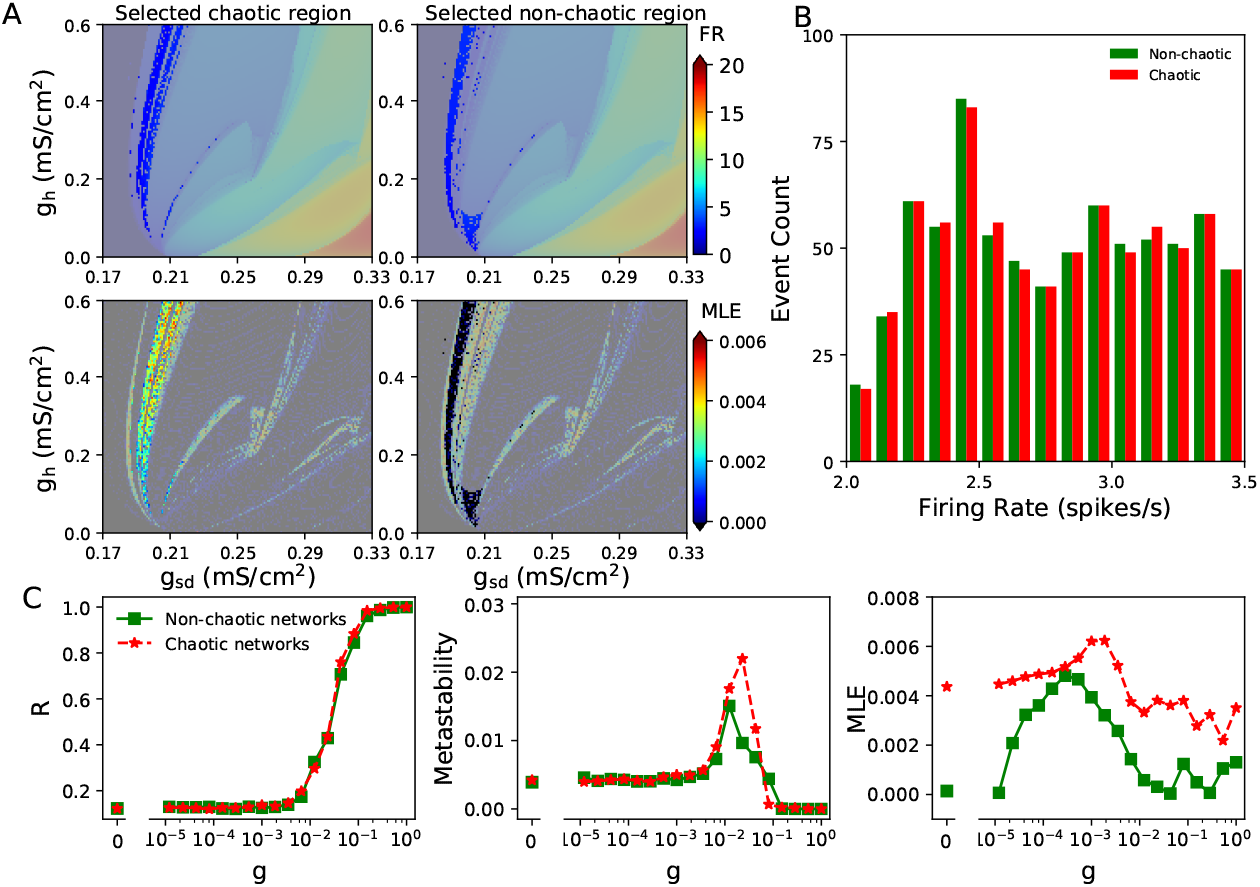
Synchronization transitions on the neural networks with parameters chosen from same distribution of firing rate of *g_sd_/g_h_* parameter space. **A**. FR (top) and MLE (bottom) obtained from each set of parameter values. The selected of blue regions in FR and MLE plot correspond to either chaotic (left) or non-chaotic oscillations (right). **B**. Histogram of firing rates in each selected parameter regions, showing the same distribution of firing rate with ranges from 2.0 to 3.5 spikes/s. **C**. Synchronization transition characterized by order parameter, metastability and the network MLE.

**S2-Fig.**
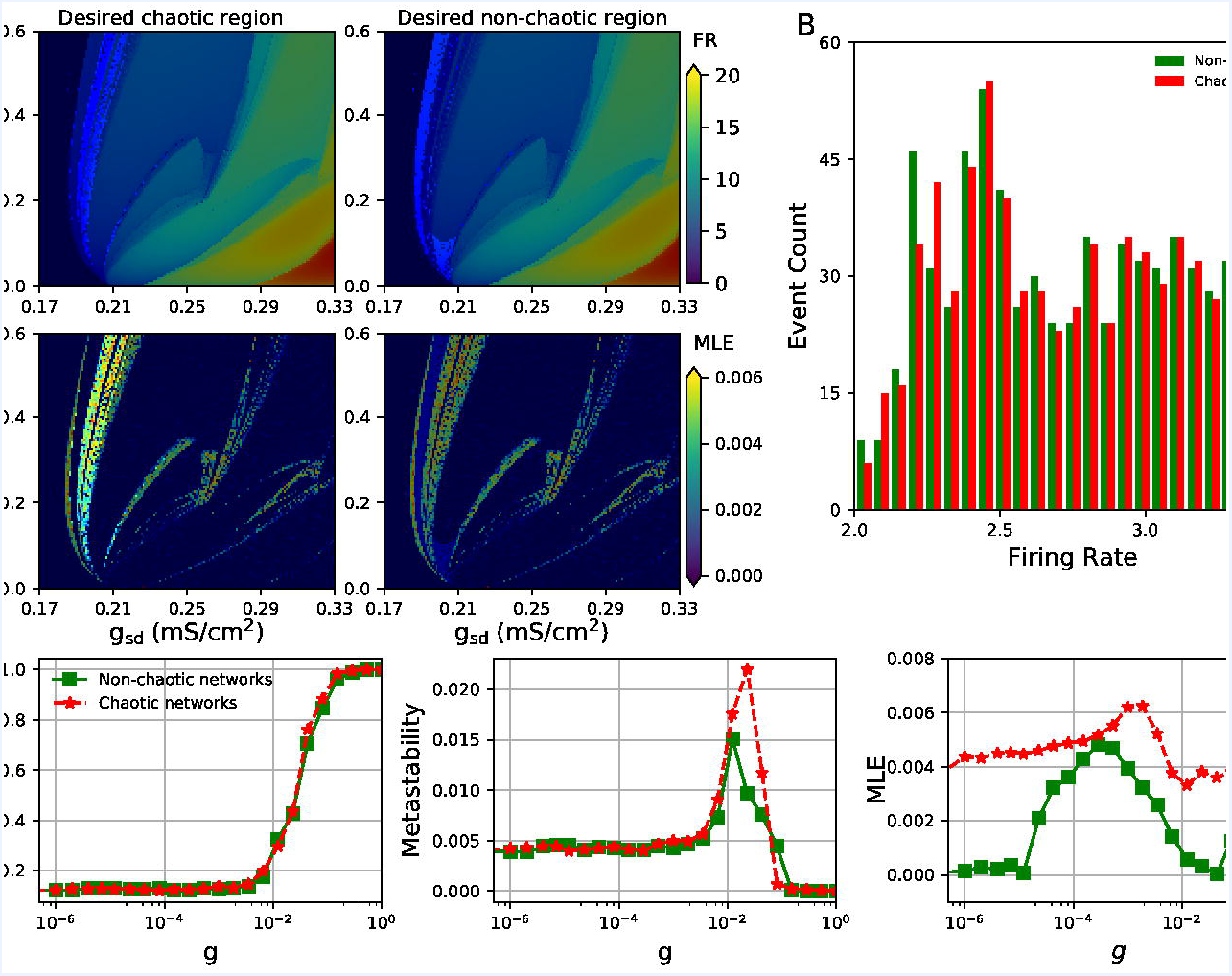
FCD in networks of chaotic and non-chaotic oscillators. **A**. FCD matrices obtained at different values of synaptic conductance *g* in networks of either chaotic or non-chaotic nodes. Below each matrix, an histogram of the values is shown. The diagonal and the neighboring values were not included. **B**. Variance of the FCD values plotted against *g*. Average of 15 simulations with different random seed for the small-world connectivity and parameters.

## Author contributions

KX and PO performed numerical simulations and analysis. JM, SC and PO performed Functional Connectivity Dynamics Analysis. KX and PO wrote the manuscript. KX, JM, SC and PO revised and approved the manuscript.

